# HydraMPP: A lightweight library for distributed massive parallel processing in Python - threading at scale

**DOI:** 10.64898/2026.06.04.730204

**Authors:** Jose L Figueroa, Richard Allen White

## Abstract

We now exist in the era of massive datasets from genomics, large language models, and all the known knowledge of humanity right at our fingertips. Much of this data is becoming more accessible; however, processing such data remains an ongoing issue across systems including high performance computing (HPC) infrastructures. Massively parallel computing (MPP) has solved this using a divide and conquer approach by splitting workloads across independent nodes (i.e., central processing units (CPU) allowing for higher scaling of data). The main engine for this in python is Ray; however, it has many issues including a large code space, security issues, debugging opacity, and memory management issues. Here, we present HydraMPP, a lightweight, ease of use and utilization, with high auditability, and with SLURM ergonomics.

## Introduction

Biological data, especially in genomics, is growing at an unprecedented rate. Unlike other big data fields, biological data comes from many different sources and is highly complex (**Stephens, 2015, Navarro et al., 2019**). This creates a need for parallel processing to analyze the data in a timely manner (**Aziz, 2021**). The complexity of the data often requires custom scripts for analysis, often written in Python, and a lack of computer science expertise in many biological laboratories further complicates how data is processed. Often, researchers writing scripts for data analysis rely on third party libraries to add functionality to their pipelines. While there are some popular Python libraries that simplify the complexities of parallel processing, they are not always a good fit for the needs of bioinformatics pipelines. They either do not provide the scalability required for large data processing across nodes on a HPC platform or they have other complexities that impede usability or ease of use. For example, joblib is a simple to use Python library that makes it trivial to convert a “for loop” to use multiple threads.

However, it does not provide the infrastructure to scale to use multiple nodes (https://github.com/joblib/joblib). Ray is another popular library that simplifies parallel computing (**Moritz et al., 2018, Karau & Lublinsky, 2022**). While Ray does provide support for multiple nodes on a HPC, it has been shown to cause conflicts when multiple users on the cluster try to use it at the same time (**Moritz et al., 2018**). It also has many dependencies that can create conflicts with other python libraries and other tools that bioinformaticians use in their pipelines.

Here we introduce HydraMPP, a Python library for massively parallel processing.

A driving force behind the development of HydraMPP is the need for a simple to use, barebones infrastructure without added complexity. HydraMPP adheres to the KISS (keep it simple and straightforward) philosophy to avoid conflicts and unnecessary overhead. This makes it easy for users with minimal scripting skills to use while not introducing overhead that may further slow down the system as the size of the job increases.

## Methods

### Software Architecture

#### Design Philosophy

HydraMPP is a library written in python (version 3 >= 3.6). Focus of the code is in simplicity of use and requiring as few dependents as possible to avoid conflicts with other libraries and improve compatibility and portability (**Fig 1**). It is designed to easily integrate into existing code without requiring massive rewrites of existing code by implementing a function “tagging” system and simple initialization. HydraMPP takes care of keeping track of available resources with the ability to automatically detect available resources. The library can also make use of multiple nodes on an HPC to further scale for large data analysis. HydraMPP is also dynamic, allowing addition of nodes after initialization (**Fig 1**).

**Figure 1:**
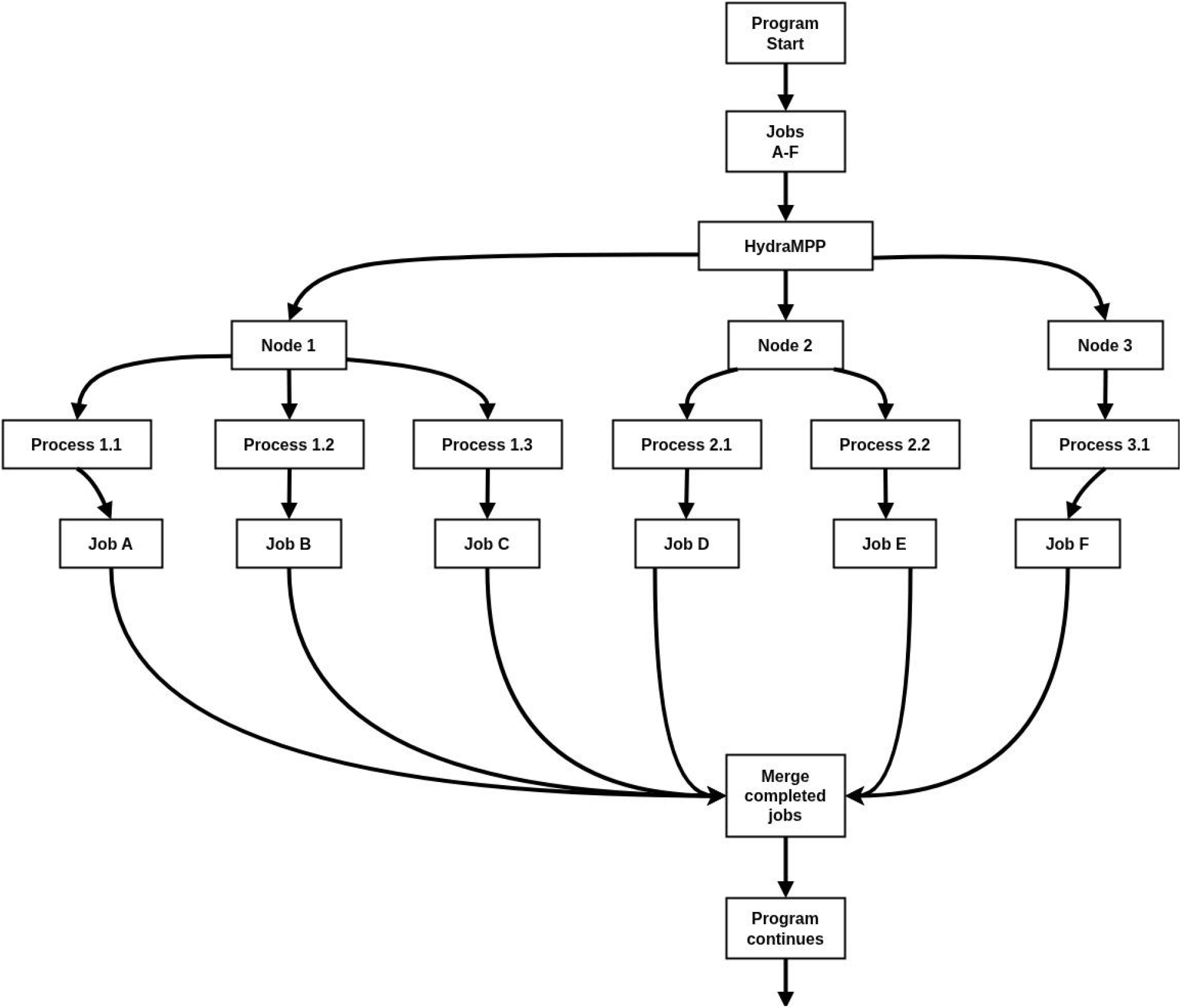
Flowgraph of HydraMPP program optimization manager. This is the general flow of how hydraMPP manages programs through distributing them across nodes via CPUs within a computing cluster.

#### Decorator-Based Task Registration

In order to make HydraMPP easy to use, Python decorators are used as tags on functions in order to register them to the library’s list of known functions, making them ready for parallel execution. This method makes it easy to implement HydraMPP on existing code without refactoring the code. This also ensures that only methods that the user explicitly wants to use in parallel processing are registered, improving security, as opposed to automatically registering undecorated functions. The user also has the option to decide how many CPUs are allocated to each method.

### Initialization and Cluster Configuration

#### Framework Initialization

Once all desired functions are tagged and other program initialization is complete, calling the init() method of HydraMPP registers the requested functions. The desired number of CPUs is also recorded, or automatically detected if not provided. At this point a list is also made with any other compute nodes on the HPC. The server is then initialized, unless running in “local” mode, and begins listening for clients. Clients are then able to connect to make their resources available. The server can be configured to allow connections for a brief time or to allow additional connections indefinitely. HydraMPP also intercepts arguments passed into the main program that begin with “--hydraMPP” for command line configuration.

#### Resource Tracking

HydraMPP automatically keeps track of available CPUs on the server along with each of the connected client nodes. It also keeps track of all running tasks, marking them as active, completed, or queued. This is updated in real time as jobs are added, running, and completed. It also has some basic error catching to identify any jobs that fail and logging error messages pointing to the cause of the error. A lightweight utility is also included for the user to check available resources and job status while HydraMPP is running.

#### SLURM Integration

HydraMPP purposefully does not automatically detect cluster environments or other running instances on HPC systems. This design consideration is to make HydraMPP non-intrusive with other running processes. Command-line arguments are used to tell HydraMPP to initialize a server/client relationship among nodes requested on SLURM systems for easy configuration. When these flags are used, the first node on the list is initialized as the server node, and it waits for the initialization and connection of the other nodes on the cluster. This makes it easy for non-computer scientists to code scripts for scaling their data analysis on HPC environments.

### Parallel Execution Model

#### Task Invocation

Registered functions tagged with the @HydraMPP.remote decorator are registered as an instance of the Worker class within HydraMPP. This class can be initialized with the default number of CPUs to allocate when running the task, or left with the default CPU count at one. The registered function can be called with the object’s remote method. Upon calling this method, the requested CPU count is evaluated and checked against the available resources. If the required resources are available, then execution starts immediately and the resource counts are adjusted accordingly. The task is tagged with a process ID and the host name of the client assigned to run the task. If the task is assigned to a connected client, the function name and arguments are encoded and sent to the client for execution. If there are not enough available resources at the time of the remote call, then the task is added to the queue. A worker thread then checks regularly for available resources and queued tasks are executed in the order that they were requested.

#### Process Spawning

HydraMPP has the option to run in different modes depending on the needs of the software. It can run in both “spawn” and “fork” modes (Linux and Mac) to balance between using shared memory space improving speed of many small jobs or efficiently leveraging Python’s processing management efficiency with low count, long running concurrent jobs. When a task is executed on the head node it is called through either a forked process or a spawned process. Forked processes have less overhead since they share memory space with the main thread, although Python does not run these threads in full parallel due to its garbage collection system. This is an efficient method for running many concurrent tasks that don’t run for too long. On the other hand, spawned processes create an entire new process creating a copy of all objects in memory space. This adds some overhead when starting new tasks, especially if there are many short running tasks. This method is best for a small number of longer running tasks. Client nodes follow the same policy as the host.

### Retrieval and Synchronization

#### Asynchronous Execution

Regardless of the method of execution, each task runs asynchronously and independently to the main program flow. This non-blocking behavior allows for more efficient use of system resources and overall reduced execution time. When a task is queued (or started) the task ID is returned to the main thread for keeping track, since the tasks are not guaranteed to return in the same order that they were executed in, they return based on when they finish running.

#### Result Collection

When tasks finish executing, HydraMPP stores the return value and sets its status as completed. The user can use the get() method to retrieve the return value. This method returns the hostname that the task ran on, the amount of time it took to run, the number of CPUs reserved for the task, whether the task is still running, and the return value. The get method requires the ID of the task requested. Tasks can be polled to check if they are done running, or the wait() method can be used to check for any completed tasks. HydraMPP.wait() returns two lists of task IDs, the completed tasks and the second list contains any pending tasks. When the get() method is called on a completed task, it removes that list from the queue. In this way, if the wait() method returns two empty lists then all tasks have completed and been retrieved.

### Resource Isolation and System-Level Design

#### Avoidance of Shared Temporary Directories

Unlike some other libraries, HydraMPP is designed to be isolated and independent of the system that it is running on. It avoids using shared temporary folders (e.g., /tmp) and does not automatically try to find any other running instances. This helps prevent interference with separate running jobs on a shared computing cluster.

This also avoids possible issues on shared systems where such shared folders can become full causing the program to crash.

#### Port and Instance Isolation

HydraMPP is also designed to be simple and use only one connection, and one port, for all communication between nodes. It is also easy to change the port number, in the case where multiple programs using HydraMPP end up sharing nodes, through the HydraMPP.init() call or using the --hydraMPP-port argument through the command line.

This helps ensure that different instances of a program or different programs using the HydraMPP library do not interfere with each other.

## Results and Discussion

### Performance Evaluation

#### Benchmarking

HydraMPP was compared against Ray, a widely used distributed computing framework for Python. Ray functions similarly to HydraMPP, although it is designed to be more resource intensive and robust. Ray utilizes MPI for distributed computing, while HydraMPP uses base Python sockets for inter-node communication. The philosophy behind this is to keep HydraMPP as easy and simple to install and light-weight as possible. We ran some performance tests using MerCat2, a fast *k*-mer counter, to compare processing time and memory usage (**Figueroa et al., 2024a**). The codebase for MerCat2 remained entirely intact except for the minimal changes to the decorators, import statements, and the return values for get() used by HydraMPP and Ray. For the comparisons, we used a random set of 100 genomes from the GTDB database (**Parks et al., 2022, Figueroa et al., 2024a**). Output files were stored on a ramdrive to avoid any issues with disk write speeds. The tests were performed first on a single node with 16 CPUs and also on a node with 36 CPUs to test the scalability. Each test was run 10 times.

#### Performance Metrics

On the node with 16 CPUs the average between HydraMPP and Ray were comparable, although Ray showed a lot more variance among the time taken to count the *k*-mers (**Fig 2**). When the available CPUs were increased to 36, hydraMPP was more efficient in using the additional resources, showing its scalability. HydraMPP was faster in both cases when using 16 CPUs or 36 CPUs due to less overhead and faster initialization for MerCat2 (**Fig 2**). HydraMPP has a lower RAM footprint when directly compared to Ray (**Fig 3**).

**Figure 2:**
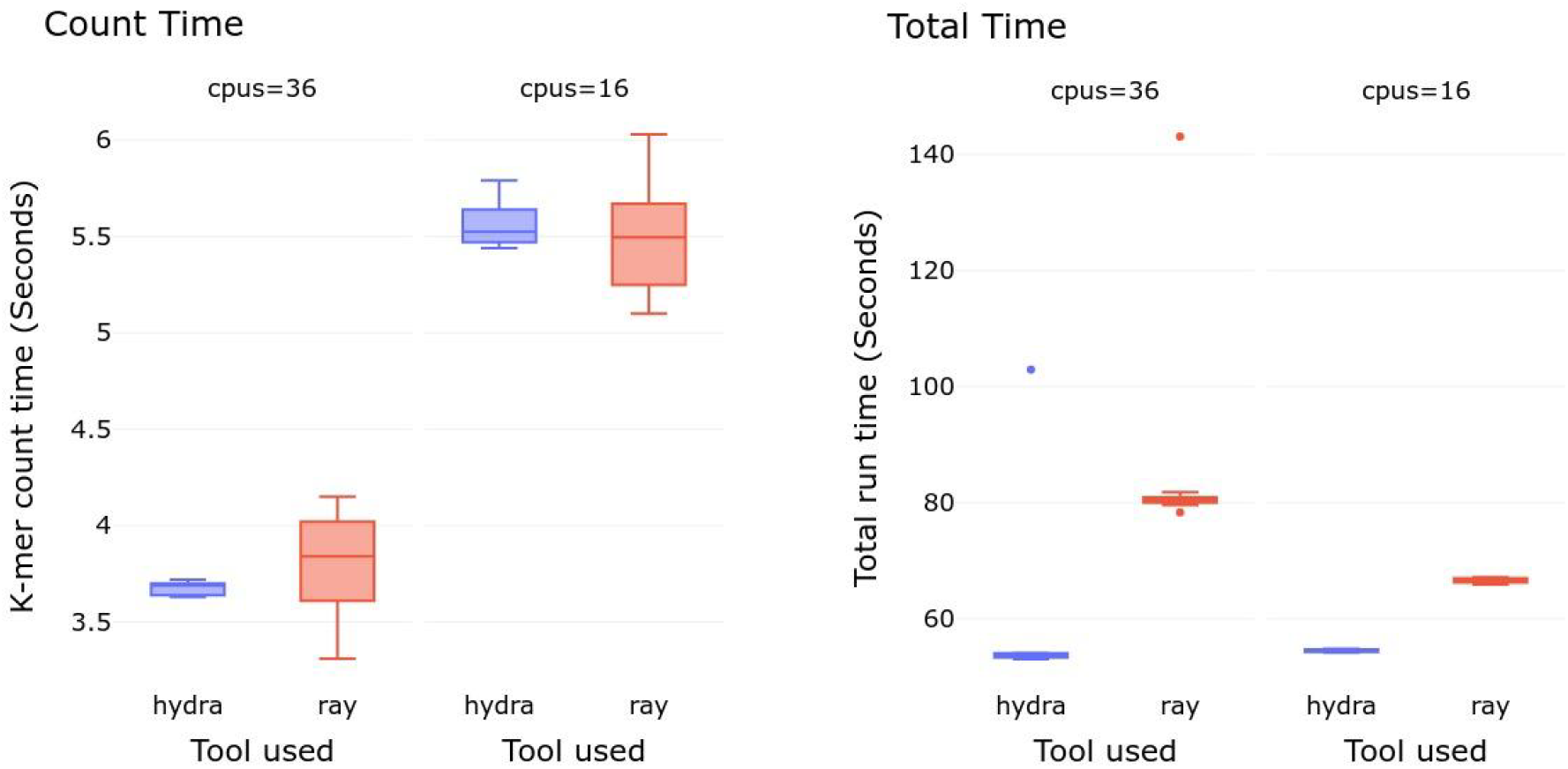
Comparison of HydraMPP vs. Ray run-time. Utilization of MerCat2 *k*-mer counter using its reference set - 100 bacteria genomes at *k* = 4 using viable numbers of CPUs.

**Figure 3:**
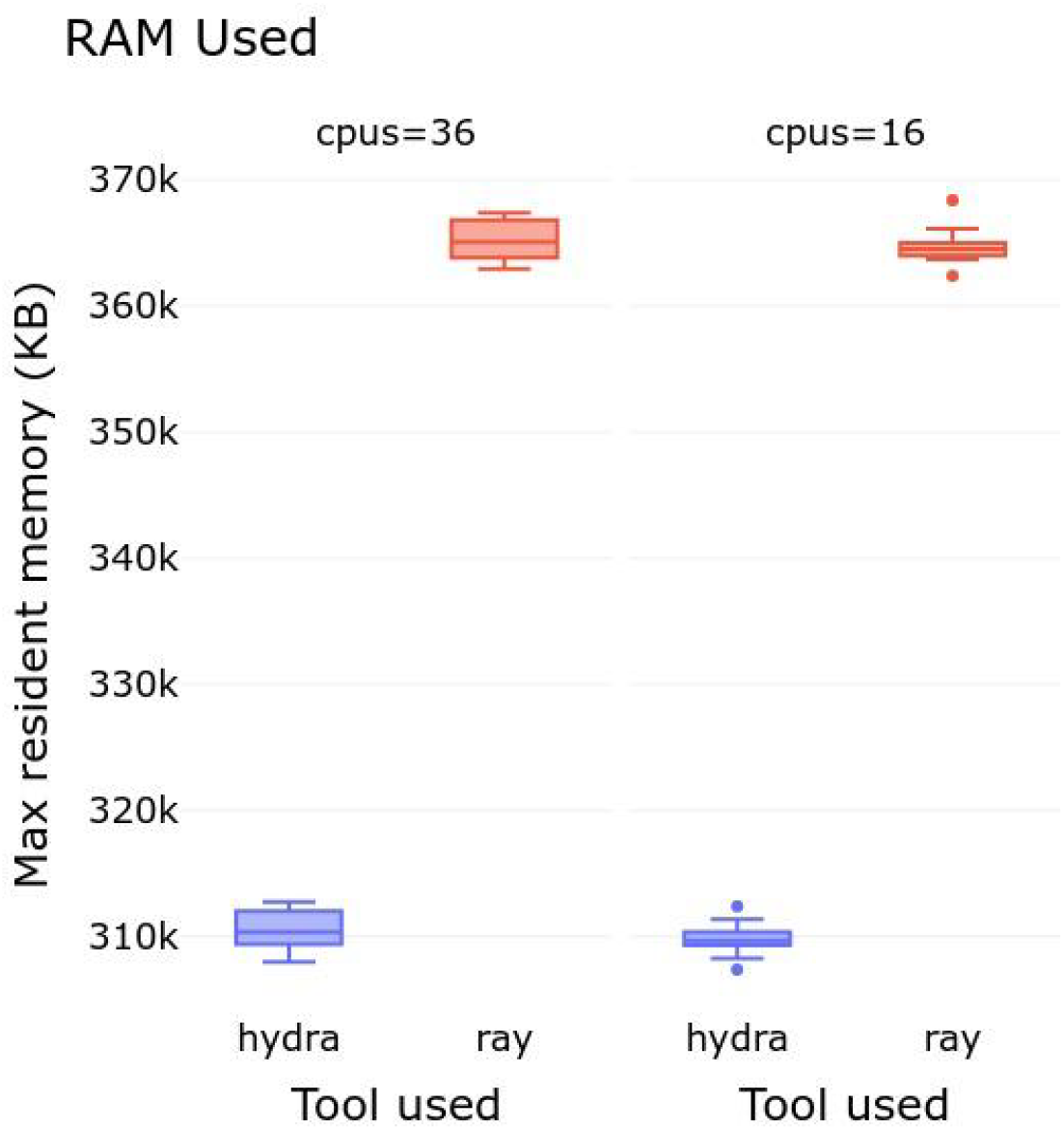
Comparison of HydraMPP vs. Ray Memory allocation. Utilization of MerCat2 *k*-mer counter using its reference set - 100 bacteria genomes at *k* = 4 using viable numbers of CPUs.

We presented that HydraMPP is an efficient and lightweight solution for parallel and distributed processing. Across the benchmarks HydraMPP was at least comparable, if not better than, Ray while also showing less variability in run time and a lower memory footprint. These results support the design philosophy of HydraMPP, that a simple framework with minimal dependencies can still provide good performance without the overhead and complexity of more feature-rich distributed platforms.

HydraMPP has been shown to function well in a few bioinformatics pipelines, including MerCat2 as discussed in the methods section for testing. It has also been successfully implemented in MetaCerberus (v1.4, **Figueroa et al., 2024b**). In deployment testing it has also been shown to be stable and reliable on computing clusters, with no conflicts when multiple users are running jobs on the cluster that utilize HydraMPP. HydraMPP shows a low runtime variance and improved scalability while maintaining a low memory footprint. These results validate the simple design philosophy of KISS, that is lightweight, and elegant software.

